# Spectral tilt underlies mathematical problem solving

**DOI:** 10.1101/601880

**Authors:** Michael J. Randazzo, Youssef Ezzyat, Michael J. Kahana

**Author notes:** Corresponding Author: Michael J. Kahana, Department of Psychology, University of Pennsylvania, 425 S. University Avenue, Philadelphia, PA 19104.

## Abstract

Neural activity associated with successful cognition appears as a tilt in the power spectrum of the local field potential, wherein increases in high-frequency power accompany decreases in low frequency power. Whereas this pattern has been shown in a wide range of memory tasks, it is unknown whether this increased spectral tilt reflects underlying memory-specific processes or rather a domain-general index of task engagement. To address the question of whether increased spectral tilt reflects increased attention to a cognitive task, we collected intracranial recordings from three hundred thirty neurosurgical patients as they performed a mathematical problem solving task. We used a mathematical problem solving task, because it allowed us to decouple task-specific processes with domain-general attention in a novel way. Using a statistical model to control for inherent problem complexity, we classified individual math problems based on whether a subject performed faster than predicted (high-attention or *fast*) or slower than predicted (low-attention, or *slow*) based on residual response times. In contrast to the domain-general attentional account, problems that took longer than predicted produced stronger evidence for the spectral tilt: widespread increases in high frequency (31–180 Hz) power and decreases in low frequency (3–17 Hz) power across frontal, temporal, and parietal cortices. The pattern emerged early within each trial and was sustained throughout the response period but was not observed in the medial temporal lobe. The data show that engaging in mathematical problem solving leads to a distributed spectral tilt pattern, even when accounting for variability in performance driven by the arithmetic demands of the problems themselves, and suggest that broadband changes in the power spectrum reflect an index of information processing in the brain beyond simple attention to the cognitive task.

## Introduction

In the domain of episodic memory, extensive prior work using both intracranial and scalp elec-troencephalography (EEG), as well as magnetoencephalography (MEG), has shown that neural activity during memory encoding exhibits broadband changes in power that correlate with memory performance (Burke, Ramayya, & Kahana, 2015). Typically, increases in high-frequency activity (HFA, >30 Hz) are associated with encoding of information that is later remembered compared to information that is later forgotten (Long, Burke, & Kahana, 2014; Burke, Long, et al., 2014; Osipova et al., 2006; Sederberg et al., 2007; Hanslmayr, Spitzer, & Bauml, 2009; Gruber, Tsivilis, Montaldi, & Mü ller, 2004). In contrast, low-frequency activity (LFA, <30 Hz) often decreases during episodic memory processing (Burke, Long, et al., 2014; Long et al., 2014; Guderian, Schott, Richardson-Klavehn, & Duzel, 2009; Staudigl & Hanslmayr, 2013; Lega, Jacobs, & Kahana, 2012; Fell, Ludowig, Rosburg, Axmacher, & Elger, 2008; Sederberg et al., 2007), although some studies have reported increases in the theta (4–8 Hz) range (Osipova et al., 2006; Hanslmayr et al., 2011; Klimesch, Doppelmayr, Russegger, & Pachinger, 1996; Burgess & Gruzelier, 2000).

The complementary increased HFA and decreased LFA (*spectral tilt* (Burke et al., 2015)) is characteristic of both memory encoding and retrieval (Burke, Sharan, et al., 2014; Kragel et al., 2017; Long et al., 2017), is observed across a range of tasks including paired associates recall (Greenberg, Burke, Haque, Kahana, & Zaghloul, 2015), and manifests in the distributed patterns of functional connectivity observed during episodic memory encoding and retrieval (Burke et al., 2013; Solomon et al., 2017). In spite of the apparent ubiquity of this broadband pattern, relatively little is known about its specificity for episodic memory processes. One interpretation is that the spectral tilt reflects engagement of contextually-mediated encoding and retrieval processes that are the hallmark of episodic memory (Tulving, 1983; Cohen & Eichenbaum, 1993). Consistent with this account, direct brain stimulation has been shown to simultaneously increase evidence for the spectral tilt and memory performance in free recall (Ezzyat et al., 2017, 2018). This account is also consistent with models proposing that HFA reflects a marker of neural information processing that can reveal with high spatial and temporal resolution the brain networks engaged in a particular cognitive task (Lachaux, Axmacher, Mormann, Halgren, & Crone, 2012; Burke et al., 2015).

However, an alternative account would propose that increased evidence for the spectral tilt could reflect a more global mechanism of orientation to the task, as opposed to specific information processing operations beyond baseline attention. Consistent with this idea, prior work has shown that attention modulates HFA (Jung et al., 2008); that task engagement compared to rest leads to a decrease in the spectral tilt in the default mode network (Miller, Weaver, & Ojemann, 2009); and that the spectral tilt is similarly increased during both memory encoding and retrieval (Kragel et al., 2017). In a typical experimental contrast comparing trials in which a subject is presumed to be engaged in the cognitive process of interest with trials in which the subject is not (e.g. *correct*/ *incorrect*), both the task-specific information processing model and the attentional model predict increased evidence for the spectral tilt. Thus, both the process-specific and attentional accounts predict that greater engagement in the cognitive task should lead to increased evidence for the spectral tilt, leaving open the question of which mechanism is more likely to drive the spectral tilt pattern.

Here, we aim to differentiate these two accounts using a mathematical problem solving task. Mathematical cognition is a skill that is included as an essential component of neuropsychological assessments and is related to a diverse array of economic, social, and psychological outcomes (Parsons & Bynner, 2005). It is also a domain in which there are inherent factors that correlate with problem difficulty and behavioral performance. For problems of mental arithmetic, factors such as the total sum and the presence of repeated digit operands are inherent to the problems themselves and affect demands on cognitive operations like executive function that are critical to task performance (Ashcraft, 1992).

To use a mathematical problem solving task to address the question of whether increased spectral tilt reflects increased attention, we collected intracranial recordings from three hundred thirty neurosurgical patients, as they performed a series of mental arithmetic problems. Taking advantage of the size of the dataset, we built a novel statistical model to account for inherent problem complexity on each trial and then classified individual problems based on whether the subject’s residual response time was faster than predicted (high-attention or *fast*) or slower than predicted (low-attention, or *slow*). After accounting for problem complexity, the attentional account would predict greater evidence for the spectral tilt for problems in which the subject performed faster than predicted by the model; in contrast, the task-related information processing account would predict increased evidence for the spectral tilt for problems in which the subject performed slower than expected (but nonetheless correctly responded). We find that difficult mathematical problem solving is associated with simultaneously increased HFA and decreased LFA, consistent with an account of the spectral tilt that is domain-general and that reflects neural information processing. We observed this pattern across broad areas of parietal, temporal, and frontal cortex, areas traditionally linked to mathematical cognition (Grabner, Ansari, et al., 2009; Daitch et al., 2016; Dehaene, Piazza, Pinel, & Cohen, 2003), but not in the hippocampus and medial temporal lobes, regions critical to the encoding and retrieval of episodic memories (Eichenbaum, 2000).

## Materials and Methods

### Participants

Three hundred thirty patients (151 females; mean age = 36 years, range 15-64 years) receiving clinical treatment for medication-resistant epilepsy were recruited to participate in this study. All patients underwent a surgical procedure in which intracranial electrodes were implanted either subdurally on the cortical surface, deep within the brain parenchyma, or both. In each case, electrode placement was determined by the clinical team. Subdural electrode contacts were arranged in strip or grid configurations with 10 mm inter-contact spacing, while depth electrodes utilized 5-10 mm inter-contact spacing. Electrophysiological data were collected as part of a multi-center collaboration at the following institutions: Dartmouth-Hitchcock Medical Center (Hanover, NH), Emory University Hospital (Atlanta, GA), Hospital of the University of Pennsylvania (Philadelphia, PA), Mayo Clinic (Rochester, MN), Thomas Jefferson University Hospital (Philadelphia, PA), Columbia University Medial Center (New York, NY), University of Texas Southwestern Medical Center (Dallas, TX), National Institutes of Health (Bethesda, MD), University of Washington Medical Center (Seattle, WA), and Freiburg University Hospital (Freiburg, Germany). The institutional review board at each institution approved the research protocol, and informed consent was obtained from the participant or the participant’s guardian.

### Experimental design

Patients participated in a mathematical problem solving task, in which they were instructed to rapidly complete a series of mental arithmetic problems. The task paradigm, developed using the Python Experiment-Programming Library (PyEPL (Geller, Schleifer, Sederberg, Jacobs, & Kahana, 2007)), was presented to participants on a laptop at the bedside, and was administered together with a delayed free recall task. The recall task involved having participants encode a list of words with subsequent recall of those words after a short delay. Participants performed the arithmetic task between the encoding and recall phases of the delayed free recall task. The memory task is not the focus of this report and will not be further discussed (Fig. 1A).

**Figure 1:**
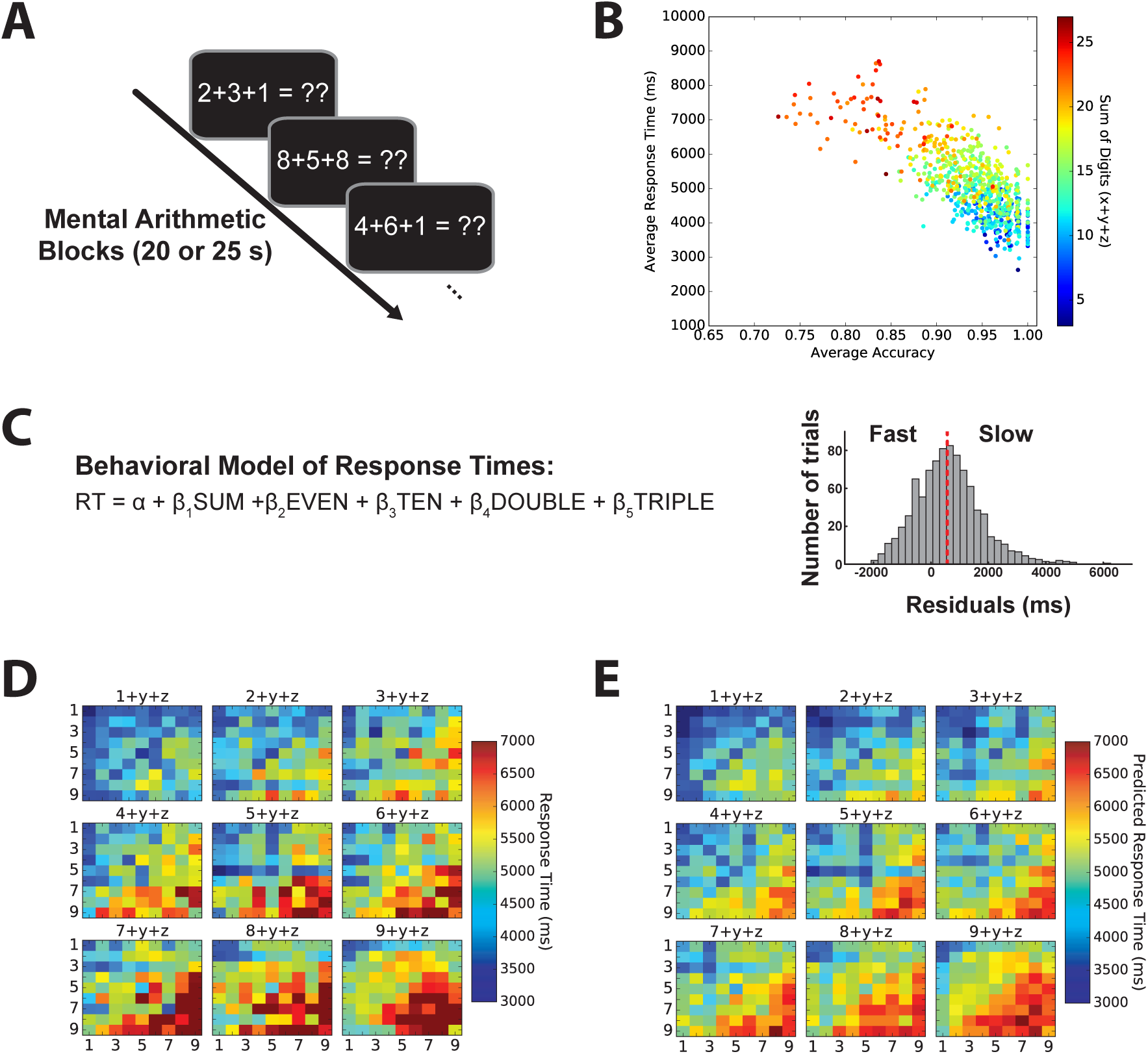
Experimental design, behavioral results, and model of arithmetic problem complexity. **A.** Participants performed blocks of a self-paced arithmetic task consisting of equations of the form of A + B + C = ??. **B.** The across-subject average accuracy and response time for each problem are graphed as a function of the problem sum. **C.** Demonstration of the method utilized for separating trials based on difficulty. *Left:* The equation used in a linear regression model of arithmetic problem complexity. *Right:* Histogram of residual response times from the behavioral model for an example subject. **D.** Average response time across subjects as a function of problem digit combination. First digit, A, is indicated above each panel, while digits B and C are represented on the *x*- and *y*-axis respectively. **E.** Predicted response times for each problem digit combination based on the aggregate subject model presented in the format of Panel D.

Each mathematical problem solving block was self-paced, which allowed participants to complete as many trials as possible; in one version of the task the interval was 20 seconds long (n = 227), while in the other the length was 25 seconds (n = 103). The interval length did not vary within-subject. On each trial, participants were presented with an arithmetic equation in the form of A + B + C = ??, where A, B, and C were randomly selected integers ranging from 1 to 9 (Fig. 1A). The participants were asked to input their answer using the numbers on the laptop keyboard and press Enter to log their response. The equation remained visible on the screen until a response was entered on the keypad, which immediately prompted the presentation of the subsequent trial. There was no limit placed on response time for a given trial, and participants were able to finish a trial once the time overall limit for the interval was reached. Each session consisted of up to 25 blocks of the arithmetic task. On average, subjects participated in two sessions (range: 1-5 sessions). We recorded accuracy and response times for each problem.

### Behavioral model

Our primary goal was to characterize the broadband changes in power that are associated with cognitively demanding mathematical problem solving, independent of the inherent complexity of the problem. To identify cognitively demanding problems, we constructed a linear regression model using aggregate subject data fit across all participants to predict their response time to each equation (Fig. 1C, left). We selected five factors for the model: (1) the sum of the digits, (2) presence of triplet digits (i.e. 3+3+3), (3) existence of any two digits with a sum of 10 (i.e. 7+3+C), (4) presence of two repeated digits (i.e. 3+3+C), and (5) sum being even or odd. These factors were chosen based on previously identified determinants of mathematical difficulty in the literature (Ashcraft, 1992) combined with distinct trends observed within our data. We also included separate confound regressors to model the mean response time for each subject. We used this model to account for baseline differences in problem difficulty in order to determine whether a participant spent more or less time solving a given problem than would be predicted by the five factors. Due to the large subject population, the same model was applied to all subjects without holding out individual subject data. We computed residual response times as the difference between a participants’ actual response time during a trial and the trial’s predicted response time; the resulting distribution of residual times for an individual subject was then separated based on the median residual, whereby *slow* (or low-attention) trials were defined as greater than the median and *fast* (or high-attention) trials were less than the median (Fig. 1C, right).

### Intracranial recordings

Intracranial EEG data were obtained at each clinical site using recording systems from a variety of manufacturers, including Bio-Logic, Blackrock, DeltaMed, Grass Tele-factor, Medtronic, Nihon-Kohden, Natus XLTek EMU128, Nicolet. Signals were sampled at 500, 512, 1000, 1024, or 2000 Hz based on the particular hardware configuration and discretion of the clinical team at each participating hospital. Recorded data were referenced to a common contact placed either intracranially, on the scalp, or on the mastoid process. A fourth order 2 Hz stop-band Butterworth notch filter was applied at 60 Hz to eliminate electrical line noise. To minimize effects from volume conduction between intracranial contacts and confounding interactions with the reference signal, a bipolar referencing montage was employed (Nunez & Srinivasan, 2006; Burke, Long, et al., 2014). Differences in signal between immediately adjacent contacts on grid, strip, and depth electrodes were calculated, creating new virtual electrodes at the midpoint between each contact pair (Burke et al., 2013).

### Anatomical localization

Anatomical localization of cortical surface (i.e. grids, strips) and depth electrodes was accomplished using independent image processing pathways. For surface electrode localization, post-implantation computed tomography (CT) images were coregistered with pre-surgical T1- or T2-weighted structural MRI scans with Advanced Normalization Tools (Avants, Epstein, Grossman, & Gee, 2008). A subset of subjects (n = 103) had post-implantation and structural scans coregistered using FMRIB’s linear image registration tool (Jenkinson, Bannister, Brady, & Smith, 2002). Individualized whole-brain cortical surfaces were then reconstructed from pre-surgical T1-weighted MRI scans using Freesurfer (Fischl et al., 2004), and electrode centroids were subsequently projected onto the cortical surface using an energy minimization algorithm (Dykstra et al., 2012). In order to cluster electrodes based on anatomical location, groups of segmented areas defined by the Desikan-Killiany atlas (Desikan et al., 2006) were designated as regions of interest (ROI). The following regions of interest were created from the specified segmented areas: superior frontal gyrus (superior frontal region), middle frontal gyrus (caudal middle frontal, rostral middle frontal regions), inferior frontal gyrus (pars opercularis, pars orbitalis, pars triangularis), inferior temporal gyrus, middle temporal gyrus, superior temporal gyrus, inferior parietal cortex (inferior parietal, supramaginal regions), superior parietal cortex (superior parietal, precuneus regions), and occipital cortex (lateral occipital region, lingual, cuneus, pericalcarine).

For localization of depth electrodes in hippocampus and medial temporal lobe (MTL), a neuroradiologist experienced in neuroanatomical localization determined each electrode’s position using post-implantation CT and MRI scans. An additional processing procedure was implemented prior to neuroradiology localization for a subset of subjects (n= 227). In this step, regions were automatically labeled on pre-implantation T2-weighted MRI scans using the automatic segmentation of hippocampal subfields (ASHS) multi-atlas segmentation method (Yushkevich et al., 2015). All cortical and subcortical regions included electrodes implanted in both hemispheres. Table 1 details the electrode coverage in each ROI across all collective subjects.

**Table 1:** Cortical and subcortical electrode coverage. This table displays the number of subjects with electrodes in a given region of interest along with the total number of electrodes recorded across all subjects. The average number of electrodes for each subject with the corresponding standard deviation is noted.

| Region of Interest (ROI) | Total Subjects | Total Electrodes | Average ± SD<br>Electrodes/Subject | Maximum<br>Electrodes/Subject |
| --- | --- | --- | --- | --- |
| Inferior Frontal Gyrus (IFG) | 228 | 1580 | $7.1 \pm 5.0$ | 27 |
| Middle Frontal Gyrus (MFG) | 220 | 2543 | $12.0 \pm 9.6$ | 51 |
| Superior Frontal Gyrus (SFG) | 144 | 1597 | $11.4 \pm 11.2$ | 53 |
| Inferior Temporal Gyrus (ITG) | 230 | 1743 | $7.8 \pm 6.1$ | 35 |
| Middle Temporal Gyrus (MTG) | 258 | 3543 | $14.3 \pm 8.8$ | 50 |
| Superior Temporal Gyrus (STG) | 244 | 2488 | $10.2 \pm 7.0$ | 28 |
| Inferior Parietal Cortex (IPC) | 230 | 2718 | $12.2 \pm 10.8$ | 60 |
| Superior Parietal Cortex (SPC) | 152 | 1176 | $7.7 \pm 7.1$ | 39 |
| Medial Temporal Lobe (MTL) | 144 | 464 | $3.2 \pm 2.1$ | 9 |
| Hippocampus (HIPPO) | 151 | 803 | $5.3 \pm 3.2$ | 17 |
| Occipital Cortex (OC) | 140 | 916 | $6.5 \pm 7.4$ | 54 |

### Spectral power

We applied the Morlet wavelet transform (wave number = 5; 8 frequencies logarithmically-spaced between 3 and 180 Hz) to all bipolar electrode EEG signals from 1,000 ms preceding math problem presentation to 1,000 ms following user input. An additional 1,000 ms buffer was included on both sides of the data segments and was subsequently discarded following the wavelet convolution to minimize edge artifacts. The resulting wavelet power estimates were then log-transformed and downsampled to 100 Hz. We normalized the resulting log-power traces using a z-transform across trials, separately within each wavelet frequency, and separately for trials within each session.

Because we were interested in examining how endogenous neural activity reflects neural information processing during *successful* mathematical problem solving, we excluded incorrect trials and trials with a response time > 30 seconds. We required a minimum of 50 such arithmetic trials to include a participant in the analysis. For the ROI analysis shown in Fig. 2A-B, continuous power traces for each subject were averaged across trials, electrodes within the ROI, and the entire response interval to yield a single power value for each trial condition (i.e. *fast, slow*), ROI, and frequency combination. This approach created a distribution of average power values across subjects in a particular region and frequency. For each ROI, we included any subject with at least one electrode localized to the ROI.

**Figure 2:**
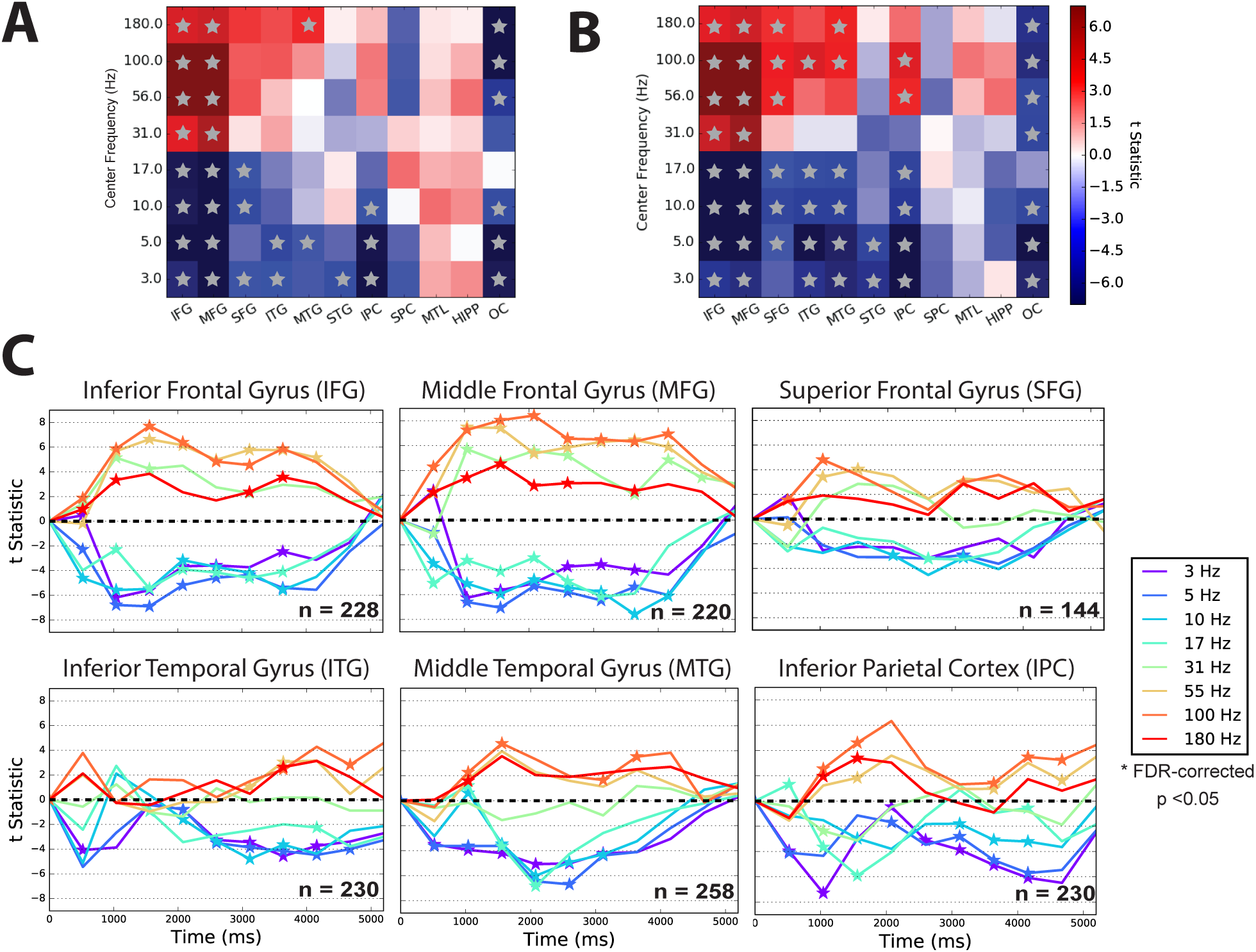
Spectral power modulation during slow and fast mental arithmetic. **A** ROI analysis contrasting spectral power from trials with longer (*slow*)response times compared to trials with shorter (*fast*) response times based on an individual’s median response time. A *t*-statistic comparing *slow* > *fast* conditions calculated for each ROI. Region and frequency pairs that exhibited an FDR-corrected difference (*q* < 0.05) between *slow* and *fast* trials are labeled with a gray star. **B** The same analysis as in (**A**); however, trials were separated with respect to the median of the *residual* response times from the behavioral model. IFG=inferior frontal gyrus; MFG=middle frontal gyrus; SFG=superior frontal gyrus; ITG=inferior temporal gyrus; MTG=middle temporal gyrus; STG= superior temporal gyrus; IPC=inferior parietal cortex; SPC=superior parietal cortex; MTL=medial temporal lobe cortex; HIPP=hippocampus. **C** Time course of spectral power changes in regions showing a spectral tilt pattern. Time along the *x*-axis represents the average post-stimulus time for each interval across all subjects. Intervals with a significant increase or decrease in spectral power (*q* < 0.05, FDR-corrected) are labeled with a star. Trials were separated by residual response times from the behavioral model as in (**B**). The number of participants included in the analysis of each ROI is shown in the lower right corner.

For analyses of the timecourse of the spectral tilt (e.g. as shown in Fig. 2C), we divided the response period for each trial into 10 non-overlapping intervals in order to account for the variable duration response times across trials. Spectral power within each interval was averaged to normalize the length of the response period, thus enabling averaging across trials. To approximate the time post-stimulus presentation that each interval represents, an average time for each interval was calculated for every subject, and the median time across subjects was displayed in lieu of the interval number. This method allowed for the characterization of broad shifts in power throughout the entire calculation process.

### Statistical analysis

We used a two-sample within-subject *t*-test to derive a measure of effect size for the comparison of spectral power between *slow* and *fast* conditions for each region and frequency. We then performed a one-sample *t*-test on the distribution of *t*-statistics across subjects to assess for the existence of a group-level difference between mathematical problem solving conditions. We used false discovery rate (FDR) to correct for multiple comparisons (Benjamini & Hochberg, 1995) with a significance level of *q* = 0.05. For Fig. 2A-B, data were corrected for all regions and frequencies, whereas for Fig. 2C, data were corrected across each time course.

## Results

### Behavioral results and model

On average, participants completed a total of 197.56 ± 12.08 (mean ± SEM) trials of the task. To assess performance on the task, we calculated each participant‘s overall accuracy (mean accuracy = 93.0 ± 6.9%). Only participants with higher than 50 percent accuracy and greater than 50 arithmetic trials were included in further analyses. 294 participants met these criteria, and therefore, 36 participants were excluded. We used response time on correct trials as our dependent measure for the behavioral model, and first sought to visualize how participant response time is affected by the total problem sum, a factor that has been previously identified as contributing to baseline problem difficulty (Ashcraft, 1992). A distinct relationship is visible, whereby increasing the total sum of digits results in longer response times and decreased accuracy (Fig 1B). This trend becomes readily apparent at larger sums, when trial combinations begin to exhibit a left upward shift of low accuracy and long response time apart from the dominant cluster with high accuracy and short response times.

Since most participants could successfully perform this task with high accuracy, we only analyzed correct trials and used response time to operationalize the information processing load required for a given problem. Fig. 1D shows the average response time for each trial combination of digits across all subjects, which illustrates the effect of problem sum in the general progression of *warm* colors (longer response times) towards the lower right corner of each subplot as well as across the entire panel, where the total sum of the digits is larger. Other patterns, such as problems in which three digits are identical (i.e. 9+9+9) or two digits sum to 10 (i.e. 5+5+C), show response times are noticeably shorter (*cool* colors) than would be predicted solely based on problem sum.

Having observed apparent relationships between arithmetic characteristics inherent to a given problem and average response times, we developed a linear regression model using aggregated subject data to predict participant response times based on properties of the problems. The model was constructed using five features of the trial equation (see Methods) that we hypothesized would be related to cognitive demand during mathematical problem solving and would therefore predict response times (Fig. 1C). Fitting the model across subjects yielded an *r*-squared value of 0.49. Normalized β-coefficients for each factor included: the sum of equation digits (*β*_1_ = 6.70), presence of an even solution (*β*_2_ = −0.06), existence of any two digits with a sum of ten (*β*_3_ = −1.15), presence of repeated digits (*β*_4_ = −0.63), and presence of triplet digits (*β*_5_ = −0.39). Using those model coefficients, we predicted response times for each possible trial equation, which are depicted in Fig. 1E. Overall, the model’s predictions exhibit similar trends to the average response times shown in Fig. 1D, which suggests that our model adequately identifies response time variability associated with measures of trial-level information processing.

### ROI analysis of spectral power changes during demanding arithmetic

We first characterized broadband changes in spectral power averaged over the response interval to determine how changes in the spectral tilt relate to variability in neural information processing during mathematical problem solving. This analysis tested the hypothesis that problems that are more cognitively demanding evoke greater evidence for the spectral tilt. We first compared trials that were above or below the median response time for each subject (Fig. 2A) before subsequently splitting trials based on whether the actual response time was above or below the predicted response time when accounting for inherent problem complexity with our behavioral model (Fig. 2B). We hypothesized that in both cases trials with longer response times were more cognitively demanding, and would therefore be associated with greater evidence for the spectral tilt. This analysis was also designed to show whether modeling inherent problem difficulty would attenuate the spectral tilt, as predicted by a process-specific account, or would lead to either no effect or an increase in evidence for the spectral tilt, as predicted by a domain-general account.

We assessed the difference in spectral power at each frequency for each electrode within subject by calculating a *t*-statistic comparing *slow* and *fast* trials; we then averaged *t*-statistics across electrodes in each ROI (across hemispheres), before assessing the effects across-subjects (one-sample *t*-test vs. 0). Low-frequency power (LFA; 3–17 Hz) during *slow* trials reliably decreased relative to *fast* trials, most prominently within the frontal cortex but also observed within areas of the temporal and parietal cortices. At the same time, the frontal lobe (including inferior and middle frontal gyri; IFG, MFG) displayed broadband increases in high frequency power (31–180 Hz); other areas including inferior temporal gyrus (ITG), middle temporal gyrus (MTG), and inferior parietal cortex (IPC) demonstrated lower magnitude increases that were not significant when correcting for multiple comparisons. The inflection point on the frequency spectrum at which the power difference shifted from negative to positive occurred between 17 and 31 Hz, consistent with previous findings observed during episodic memory (Burke, Long, et al., 2014). The IFG and MFG regions showed strongest evidence for the spectral tilt, consistent with a role in domain-general manipulation and organization of information in working memory (Owen et al., 1998; Blumenfeld & Ranganath, 2006; Kong et al., 2005; Ischebeck, Zamarian, Egger, Schocke, & Delazer, 2007). In contrast, the occipital cortex (OC), which is responsible for similar visual processing during both trial types, does not exhibit a spectral tilt.

We next reclassified trials based on our behavioral model of intrinsic mathematical problem difficulty, to determine whether controlling for problem-level complexity would eliminate the spectral tilt pattern we observed when using raw response time to bin trials. We used the model to predict response times for each trial and then split trials into *slow* and *fast* conditions based on the residuals (see Methods). Using the same analysis from Fig. 2A, we found that controlling for trial-level variability led to a stronger spectral tilt, in contrast to the prediction of the process specific model and consistent with a domain general account (Fig. 2B). ITG, MTG, and IPC all showed significant low-frequency power decreases between 3–17 Hz, along with significant high-frequency power increases from 56–180 Hz. The increase in high-frequency power within the superior frontal gyrus (SFG) also reached significance at all frequencies between 56–180 Hz. Furthermore, the decrease in low-frequency power was more widespread and encompassed all ROIs including the hippocampus and medial temporal lobes.

### Timecourse of spectral power changes in arithmetic

Having characterized the aggregate pattern of neural activity across the brain during cognitively demanding problem solving, we next investigated the temporal dynamics of the spectral tilt across the response period. To align trials with varying response times, we first performed a vincentization of the response period, whereby the response period for each trial was divided into 10 intervals and average power computed within each interval. This allows us to statistically compare (across trials and subjects) intervals that were matched for their relative within-response period position. Fig. 2C shows the average time course of activity in regions that demonstrated a significant decrease in LFA combined with a significant increase in HFA in Fig. 2B. The most significant response was observed in frontal lobe ROIs (IFG and MFG), where all of the high-frequencies (*warm* colors) exhibited significantly increased power while the low-frequencies (*cool* colors) exhibited significantly decreased power that persisted from stimulus presentation to subject response.

In the temporal lobes, the ITG showed a late increase in high-frequency power and reduction in low-frequency power compared to the MTG, which showed two high-frequency power peaks and a decrease in low-frequency power that was sustained for much of the response period. In contrast, IPC showed an initial high-frequency peak in the first half of the response period along with significantly reduced low-frequency power. Taken together, these data suggest that cognitive demand modulates the spectral tilt most strongly in frontal regions in a way that is consistent across the response period, suggesting sustained engagement of neural activity in these areas during *difficult* mathematical problem solving.

## Discussion

We evaluated the link between the spectral tilt and cognitive demand in the context of a mathematical problem solving task by recording intracranial EEG from cortical and subcortical electrodes implanted in a large sample of neurosurgical patients. By analyzing a large dataset to achieve extensive electrode coverage, we could evaluate whole-brain spectral dynamics during mathematical problem solving. After controlling for problem difficulty, problems that subjects answered correctly but slower than predicted by the model demonstrated greater evidence for the spectral tilt, most strongly in areas of the lateral frontal lobe. The data show that neural information processing during mathematical problem solving exhibits similar biomarkers of successful performance as found in other domains, such as episodic memory encoding and retrieval, in a way that is inconsistent with an attention-based account of the spectral tilt. The data suggest that the spectral tilt may reflect the presence of *desirable difficulties* that reflect states of information processing associated with successful cognition.

### Electrophysiological and cognitive basis of the spectral tilt

There is substantial evidence that broadband high-frequency power in the local field potential can be used to index unit firing of the underlying neural population (Lachaux et al., 2012; Merker, 2013) that is correlated with the blood-oxygen level dependent (BOLD) fMRI response (Conner, Ellmore, Pieters, DiSano, & Tandon, 2011; Winawer et al., 2013) and multi-unit activity (Manning, Jacobs, Fried, & Kahana, 2009). For example, high-frequency power has been linked to information processing across several cognitive domains including sensorimotor integration (Crone, Sinai, & Korzeniewska, 2006; Cheyne, Bells, Ferrari, Gaetz, & Bostan, 2008; Miller et al., 2007; Crone, Miglioretti, Gordon, & Lesser, 1998), auditory speech perception (Chang et al., 2011), visual recognition (Hermes, Miller, Wandell, & Winawer, 2015), and memory encoding and retrieval (Foster, Dastjerdi, & Parvizi, 2012; Burke, Long, et al., 2014; Howard et al., 2003). Previous studies that used iEEG to study mental arithmetic also measured high-frequency power in order to detect calculation-specific activity (Ueda, Brown, Kojima, Juhász, & Asano, 2015; Hermes, Rangarajan, et al., 2015; Daitch et al., 2016). In addition to a strong relationship between high-frequency power and neural activity, prior work has also observed a concurrent reduction in low frequency activity (Ezzyat et al., 2017; Burke, Long, et al., 2014; Burke, Sharan, et al., 2014; Greenberg et al., 2015; Long et al., 2014). Unlike the largely asynchronous high-frequency power modulations, these low frequency power changes undergo synchronization, which may provide a mechanism for inter-regional communication (Burke et al., 2013; Solomon et al., 2017).

Our findings demonstrate that the broadband changes in spectral power previously observed across multiple cognitive domains are also present during periods of cognitively demanding mathematical problem solving. Because our analysis focused exclusively on correct trials, our results are unlikely to be related to fluctuations in attention to the task that could sometimes lead to incorrect responses. Instead, trials that required longer processing time showed greater evidence for the spectral tilt, an effect that was not driven by arithmetic properties of the problems themselves that are known to correlate with response time. The data suggest that cognitively demanding mathematical problem solving exhibits a pattern of whole-brain broadband spectral power that is similar to that observed during periods of successful episodic memory formation and retrieval (Kragel et al., 2017).

An important direction for future work will be to directly compare the neural biomarkers of success across cognitive domains, for example mathematical problem solving and episodic memory encoding/retrieval. Although it was not the focus of this manuscript, it is interesting to note the qualitative similarities between the whole-brain spectral tilt during cognitively demanding mathematical problem solving and periods of successful memory encoding and retrieval. The consistency in neural activity between difficult mathematical problem solving and episodic memory processes is consistent with the notion of *desirable difficulties* in memory, whereby engaging cognitively demanding learning leads to better long-term memory retention (BjorK & Kroll, 2015; Karpicke & Roediger, 2008). One possibility is that the similar patterns of broadband spectral modulation reflect a state of neural information processing that is associated with periods of successful cognition (Hasson, Chen, & Honey, 2015).

### A behavioral model of mathematical problem solving

We introduced a novel behavioral model to account for trial-level variability in arithmetic factors that are known to correlate with difficulty and response time (Ashcraft, 1992). Previous studies have generally defined levels of arithmetic difficulty by pre-selecting trials based on intrinsic features related to equation difficulty or by having participants perform pre-experimental training to selectively reduce the difficulty of trained equations (Grabner, Ischebeck, et al., 2009; Ischebeck, Zamarian, Schocke, & Delazer, 2009; Ischebeck et al., 2007). The most common intrinsic features designed to raise procedural complexity included increasing the magnitude of the digits, for example from single-digit to double-digit (Vansteensel et al., 2014; Grabner, Ansari, et al., 2009; Ueda et al., 2015), or choosing problems that require performing carrying or borrowing (Kong et al., 2005; Klein et al., 2010). Our approach is distinct from these earlier studies because we sought to explicitly account for and remove the influence of arithmetic factors on response times. We then used the resulting model residuals to bin trials based on residual response times, thus identifying spectral signatures associated with endogenous variability in a person’s cognitive state (Gilden, Thornton, & Mallon, 1995).

### Whole-brain contributions to mathematical problem solving

By using the spectral tilt to index neural information processing and analyzing intracranial EEG recordings in a large dataset, our study was able to replicate and extend to the whole-brain level previous studies that have identified core mechanisms of mathematical problem solving in specific neural populations (Dastjerdi, Ozker, Foster, Rangarajan, & Parvizi, 2013; Daitch et al., 2016; Ueda et al., 2015; Vansteensel et al., 2014). Our findings demonstrate that regions of the frontal cortex remain activated throughout the response interval with a spectral difference arising shortly after cue presentation and normalizing immediately before response production (Fig. 2C). Previous fMRI studies have shown modulations of BOLD signal in IFG in response to manipulations of equation complexity and level of practice with specific arithmetic problems (Kazui, Kitagaki, & Mori, 2000; Kong et al., 2005; Delazer et al., 2003; Arsalidou & Taylor, 2011). Yet, prior intracranial EEG studies have failed to identify significant high frequency activity within this region. Our results are therefore consistent with the fMRI literature and demonstrate a plausible temporal course of activation, whereby equation presentation causes an initial peak in cognitive demand followed by persistent activity until a solution is obtained.

Our findings in the parietal cortex are consistent with the prominent Triple Code model (Dehaene et al., 2003) of mathematical cognition, as well as findings from intracranial EEG. The Triple Code model predicts the existence of three numerical representations in parietal cortex defined by the angular gyrus, intraparietal sulcus, and superior parietal system. The angular gyrus, for example, has been linked to arithmetic fact retrieval and learning (Ischebeck et al., 2009; Klein, Moeller, Glauche, Weiller, & Willmes, 2013; Grabner et al., 2007), while the intraparietal sulcus has a role in representing numerical quantities (Kadosh & Walsh, 2009). These regions, which are part of our IPC ROI, have demonstrated calculation-related high-frequency power in several intracranial EEG studies (Dastjerdi et al., 2013; Daitch et al., 2016; Ueda et al., 2015; Vansteensel et al., 2014) that is also related to problem difficulty (Vansteensel et al., 2014).

Prior work has also identified modulations in functional coupling between the IPC and inferior temporal lobe during distinct stages of numerical processing (Daitch et al., 2016). Our work extends these findings by showing a robust spectral tilt response in ITG that is emphasized when separating trials based on cognitive demand (Fig. 2A-B). When examining the timing of this activity, a predominantly late response was detected, which aligns with the idea that ITG participates in direct computation in addition to early visual numeral encoding. The contribution of MTG to mental calculation has been less clearly elucidated in the prior literature. Lesions in this area cause deficits in rote recall of arithmetic facts (Dehaene & Cohen, 1997), while fMRI functional connectivity increases during easier arithmetic (Klein et al., 2013, 2016). It may seem surprising then that we observe a significant spectral tilt in this area during cognitively demanding mathematical problem solving. Two possible explanations are that MTG is recruited during situations of arithmetic fact retrieval as well as cognitively demanding arithmetic computation, or that some element of arithmetic fact retrieval contributes to performance during cognitively demanding problem solving. Future work will be necessary to adjudicate between these possibilities.

## Conclusion

We recorded intracranial EEG in a large sample of participants to obtain extensive cortical and subcortical electrode coverage, with which we characterized whole-brain patterns of neural activity during cognitively demanding mathematical problem solving. Spectral analysis revealed a widespread spectral tilt pattern characterized by increased high-frequency power and decreased low-frequency power. This broadband pattern was present across frontal, parietal and temporal cortical areas for problems that required high levels of information processing, a pattern similar to that observed in previous studies in other cognitive domains such as episodic memory. The data suggest that broadband shifts in the power spectrum of neural activity arise from task-related information processing and are unlikely to reflect basic attention or orientation to the task.

## Acknowledgements

We thank Blackrock Microsystems for providing neural recording and stimulation equipment. This work was supported by the DARPA Restoring Active Memory (RAM) program (Cooperative Agreement N66001-14-2-4032). The views, opinions, and/or findings contained in this material are those of the authors and should not be interpreted as representing the official views or policies of the Department of Defense or the U.S. Government.

## References

Arsalidou, M., & Taylor, M. J. (2011). Is 2+2=4? Meta-analyses of brain areas needed for numbers and calculations. NeuroImage, 54(3), 2382–2393. Retrieved from http://dx.doi.org/10.1016/j.neuroimage.2010.10.009 doi:10.1016/j.neuroimage.2010.10.009

Ashcraft, M. H. (1992). Cognitive arithmetic: A review of data and theory. Cognition, 44(1-2), 75–106.

Avants, B. B., Epstein, C. L., Grossman, M., & Gee, J. C. (2008). Symmetric diffeomorphic image registration with cross-correlation: evaluating automated labeling of elderly and neurodegenerative brain. Medical Image Analysis, 12(1), 26–41.

Benjamini, Y., & Hochberg, Y. (1995). Controlling the False Discovery Rate: a practical and powerful approach to multiple testing. Journal of Royal Statistical Society, Series B, 57, 289–300.

BjorK, R. A., & Kroll, J. F. (2015). Desirable difficulties in vocabulary learning. The American journal of psychology, 128(2), 241.

Blumenfeld, R., & Ranganath, C. (2006). Dorsolateral prefrontal cortex promotes long-term memory formation through its role in working memory organization. The Journal of neuroscience, 26(3), 916–925.

Burgess, A. P., & Gruzelier, J. H. (2000). Short duration power changes in the EEG during recognition memory for words and faces. Psychophysiology, 37, 596–606.

Burke, J. F., Long, N. M., Zaghloul, K. A., Sharan, A. D., Sperling, M. R., & Kahana, M. J. (2014). Human intracranial high-frequency activity maps episodic memory formation in space and time. NeuroImage, 85 Pt. 2, 834–843. doi:0.1016/j.neuroimage.2013.06.067

Burke, J. F., Ramayya, A. G., & Kahana, M. J. (2015). Human intracranial high-frequency activity during memory processing: Neural oscillations or stochastic volatility? Current Opinion in Neurobiology, 31, 104–110.

Burke, J. F., Sharan, A. D., Sperling, M. R., Ramayya, A. G., Evans, J. J., Healey, M. K., … Kahana, M. J. (2014). Theta and high–frequency activity mark spontaneous recall of episodic memories. Journal of Neuroscience, 34(34), 11355–11365. doi:10.1523/JNEUROSCI.2654-13.2014

Burke, J. F., Zaghloul, K. A., Jacobs, J., Williams, R. B., Sperling, M. R., Sharan, A. D., & Kahana, M. J. (2013). Synchronous and asynchronous theta and gamma activity during episodic memory formation. Journal of Neuroscience, 33(1), 292–304. doi:10.1523/JNEUROSCI.2057-12.2013

Chang, E. F., Edwards, E., Nagarajan, S. S., Fogelson, N., Dalal, S. S., Canolty, R. T., … Knight, R. T. (2011). Cortical spatio-temporal dynamics underlying phonological target detection in humans. Journal of cognitive neuroscience, 23(6), 1437–1446.

Cheyne, D., Bells, S., Ferrari, P., Gaetz, W., & Bostan, A. C. (2008). Self-paced movements induce high-frequency gamma oscillations in primary motor cortex. Neuroimage, 42(1), 332–342.

Cohen, N. J., & Eichenbaum, H. (1993). Memory, amnesia, and the hippocampal system. Cambridge, MA: MIT.

Conner, C. R., Ellmore, T. M., Pieters, T. A., DiSano, M. A., & Tandon, N. (2011, Sep). Variability of the relationship between electrophysiology and bold-fmri across cortical regions in humans. J Neurosci, 31(36), 12855–65. doi:10.1523/JNEUROSCI.1457-11.2011

Crone, N. E., Miglioretti, D. L., Gordon, B., & Lesser, R. P. (1998). Functional mapping of human sensorimotor cortex with electrocorticographic spectral analysis. II. Event-related synchronization in the gamma band. Brain, 121(12), 2301–2315.

Crone, N. E., Sinai, A., & Korzeniewska, A. (2006). High-frequency gamma oscillations and human brain mapping with electrocorticography. Progress in Brain Research, 159, 275–295.

Daitch, A. L., Foster, B. L., Schrouff, J., Rangarajan, V., Kaşikçi, I., Gattas, S., & Parvizi, J. (2016). Mapping human temporal and parietal neuronal population activity and functional coupling during mathematical cognition. Proceedings of the National Academy of Sciences, 201608434. Retrieved from http://www.pnas.org/lookup/doi/10.1073/pnas.1608434113 doi:1.1073/pnas.1608434113

Dastjerdi, M., Ozker, M., Foster, B. L., Rangarajan, V., & Parvizi, J. (2013). Numerical processing in the human parietal cortex during experimental and natural conditions. Nature communications, 4, 2528. Retrieved from http://www.pubmedcentral.nih.gov/articlerender.fcgi?artid=3826627{\&}tool=pmcentrez{\&}rendertype=abstract doi:10.1038/ncomms3528

Dehaene, S., & Cohen, L. (1997). Cerebral pathways for calculation: Double dissociation between rote verbal and quantitative knowledge of arithmetic. Cortex, 33(2), 219–250.

Dehaene, S., Piazza, M., Pinel, P., & Cohen, L. (2003). Three parietal circuits for number processing. Cognitive neuropsychology, 20(3), 487–506. Retrieved from http://www.ncbi.nlm.nih.gov/pubmed/20957581{\%}5Cnhttp://www.scopus.com/inward/record.url?eid=2-s2.0-0013535126{\&}partnerID=tZOtx3y1 doi:10.1080/02643290244000239

Delazer, M., Domahs, F., Bartha, L., Brenneis, C., Lochy, A., Trieb, T., & Benke, T. (2003). Learning complex arithmetic - An fMRI study. Cognitive Brain Research, 18(1), 76–88. doi:10.1016/j.cogbrainres.2003.09.005

Desikan, R., Segonne, B., Fischl, B., Quinn, B., Dickerson, B., Blacker, D., … Killiany, N. (2006). An automated labeling system for subdividing the human cerebral cortex on MRI scans into gyral based regions of interest. NeuroImage, 31(3), 968–80.

Dykstra, A. R., Chan, A. M., Quinn, B. T., Zepeda, R., Keller, C. J., Cormier, J., … Cash, S. S. (2012). Individualized localization and cortical surface-based registration of intracranial electrodes. Neuroimage, 59(4), 3563–3570.

Eichenbaum, H. (2000, Oct). A cortical-hippocampal system for declarative memory. Nature Reviews. Neuroscience, 1(1), 41–50.

Ezzyat, Y., Kragel, J. E., Burke, J. F., Levy, D. F., Lyalenko, A., Wanda, P., … Kahana, M. J. (2017). Direct brain stimulation modulates encoding states and memory performance in humans. Current Biology, 27(9), 1251–1258.

Ezzyat, Y., Wanda, P., Levy, D., Kadel, A., Aka, A., Pedisich, I., … Kahana, M. (2018). Closed-loop stimulation of temporal cortex rescues functional networks and improves memory. Nature Communications, 9(1), 365. doi:10.1038/s41467-017-02753-0

Fell, J., Ludowig, E., Rosburg, T., Axmacher, N., & Elger, C. (2008). Phase-locking within human mediotemporal lobe predicts memory formation. Neuroimage, 43(2), 410–419.

Fischl, B., van der Kouwe, A., Destrieux, C., Halgren, E., Ségonne, F., Salat, D. H., … others (2004). Automatically parcellating the human cerebral cortex. Cerebral Cortex, 14(1), 11–22.

Foster, B. L., Dastjerdi, M., & Parvizi, J. (2012). Neural populations in human posteromedial cortex display opposing responses during memory and numerical processing. Proceedings of the National Academy of Sciences, 109(38), 15514–15519. doi:10.1073/pnas.1206580109

Geller, A. S., Schleifer, I. K., Sederberg, P. B., Jacobs, J., & Kahana, M. J. (2007). PyEPL: A cross-platform experiment-programming library. Behavior Research Methods, 39(4), 950–958.

Gilden, D. L., Thornton, T., & Mallon, M. W. (1995). 1/f noise in human cognition. Science, 267(5205), 1837–1839.

Grabner, R. H., Ansari, D., Koschutnig, K., Reishofer, G., Ebner, F., & Neuper, C. (2009). To retrieve or to calculate? Left angular gyrus mediates the retrieval of arithmetic facts during problem solving. Neuropsychologia, 47(2), 604–608. doi:10.1016/j.neuropsychologia.2008.10.013

Grabner, R. H., Ansari, D., Reishofer, G., Stern, E., Ebner, F., & Neuper, C. (2007). Individual differences in mathematical competence predict parietal brain activation during mental calculation. Neuroimage, 38(2), 346–356.

Grabner, R. H., Ischebeck, A., Reishofer, G., Koschutnig, K., Delazer, M., Ebner, F., & Neuper, C. (2009). Fact learning in complex arithmetic and figural-spatial tasks: The role of the angular gyrus and its relation to mathematical competence. Human Brain Mapping, 30(9), 2936–2952. doi:10.1002/hbm.20720

Greenberg, J. A., Burke, J. F., Haque, R., Kahana, M. J., & Zaghloul, K. A. (2015, Jul). Decreases in theta and increases in high frequency activity underlie associative memory encoding. Neuroimage, 114, 257–263. doi:10.1016/j.neuroimage.2015.03.077

Gruber, T., Tsivilis, D., Montaldi, D., & Müller, M. (2004). Induced gamma band responses: An early marker of memory encoding and retrieval. Neuroreport, 15, 1837–1841.

Guderian, S., Schott, B., Richardson-Klavehn, A., & Duzel, E. (2009). Medial temporal theta state before an event predicts episodic encoding success in humans. Proceedings of the National Academy of Sciences, 106(13), 5365.

Hanslmayr, S., Spitzer, B., & Bauml, K. (2009). Brain oscillations dissociate between semantic and nonsemantic encoding of episodic memories. Cerebral Cortex, 19(7), 1631–1640.

Hanslmayr, S., Volberg, G., Wimber, M., Raabe, M., Greenlee, M. W., & Bäumel, K. H. T. (2011). The relationship between brain oscillations and bold signal during memory formation: A combined eeg-fmri study. Journal of Neuroscience, 31(44), 15674–15680.

Hasson, U., Chen, J., & Honey, C. J. (2015). Hierarchical process memory: memory as an integral component of information processing. Trends in cognitive sciences, 19(6), 304–313.

Hermes, D., Miller, K. J., Wandell, B. A., & Winawer, J. (2015). Gamma oscillations in visual cortex: the stimulus matters. Trends in cognitive sciences, 19(2), 57–58.

Hermes, D., Rangarajan, V., Foster, B. L., King, J.-R., Kasikci, I., Miller, K. J., & Parvizi, J. (2015). Electrophysiological Responses in the Ventral Temporal Cortex During Reading of Numerals and Calculation. Cerebral Cortex, 27(1), bhv250. Retrieved from http://www.cercor.oxfordjournals.org/lookup/doi/10.1093/cercor/bhv250 xdoi:10.1093/cercor/bhv250

Howard, M. W., Rizzuto, D. S., Caplan, J. C., Madsen, J. R., Lisman, J., Aschenbrenner-Scheibe, R., … Kahana, M. J. (2003). Gamma oscillations correlate with working memory load in humans. Cerebral Cortex, 13, 1369–1374.

Ischebeck, A., Zamarian, L., Egger, K., Schocke, M., & Delazer, M. (2007). Imaging early practice effects in arithmetic. NeuroImage, 36(3), 993–1003. Retrieved from http://dx.doi.org/10.1016/j.neuroimage.2007.03.051 doi:10.1016/j.neuroimage.2007.03.051

Ischebeck, A., Zamarian, L., Schocke, M., & Delazer, M. (2009). Flexible transfer of knowledge in mental arithmetic - An fMRI study. NeuroImage, 44(3), 1103–1112. Retrieved from http://dx.doi.org/10.1016/j.neuroimage.2008.10.025 doi:10.1016/j.neuroimage.2008.10.025

Jenkinson, M., Bannister, P., Brady, M., & Smith, S. (2002). Improved optimisation for the robust and accurate linear registration and motion correction of brain images. NeuroImage, 17(2), 825–841.

Jung, J., Mainy, N., Kahane, P., Minotti, L., Hoffmann, D., Bertrand, O., & Lachaux, J. (2008). The neural bases of attentive reading. Human Brain Mapping, 29(1193–1206).

Kadosh, R. C., & Walsh, V. (2009). Numerical representation in the parietal lobes: Abstract or not abstract? Behavioral and brain sciences, 32(3-4), 313–328.

Karpicke, J. D., & Roediger, H. L., III. (2008, February). The critical importance of retrieval for learning. Science, 319, 966–968.

Kazui, H., Kitagaki, H., & Mori, E. (2000). Cortical activation during retrieval of arithmetical facts and actual calculation: A functional magnetic resonance imaging study. Psychiatry and Clinical Neurosciences, 54(4), 479–485. doi:10.1046/j.1440-1819.2000.00739.x

Klein, E., Moeller, K., Glauche, V., Weiller, C., & Willmes, K. (2013). Processing Pathways in Mental Arithmetic-Evidence from Probabilistic Fiber Tracking. PLoS ONE, 8(1). doi:10.1371/journal.pone.0055455

Klein, E., Suchan, J., Moeller, K., Karnath, H. O., Knops, A., Wood, G., … Willmes, K. (2016). Considering structural connectivity in the triple code model of numerical cognition: differential connectivity for magnitude processing and arithmetic facts. Brain Structure and Function, 221(2), 979–995. doi:10.1007/s00429-014-0951-1

Klein, E., Willmes, K., Dressel, K., Domahs, F., Wood, G., & Nuerk, H.-c. (2010). Categorical and continuous - disentangling the neural correlates of the carry effect in multi-digit addition. Behavioral and Brain Functions, 6(1), 70. Retrieved from http://www.behavioralandbrainfunctions.com/content/6/1/70 xdoi:10.1186/1744-9081-6-70

Klimesch, W., Doppelmayr, M., Russegger, H., & Pachinger, T. (1996). Theta band power in the human scalp EEG and the encoding of new information. NeuroReport, 7, 1235–1240.

Kong, J., Wang, C., Kwong, K., Vangel, M., Chua, E., & Gollub, R. (2005). The neural substrate of arithmetic operations and procedure complexity. Cognitive Brain Research, 22(3), 397–405. doi:10.1016/j.cogbrainres.2004.09.011

Kragel, J. E., Ezzyat, Y., Sperling, M. R., Gorniak, R., Worrell, G. A., Berry, B. M., … Kahana, M. J. (2017, July). Similar patterns of neural activity predict memory function during encoding and retrieval. NeuroImage, 155, 60–71.

Lachaux, J. P., Axmacher, N., Mormann, F., Halgren, E., & Crone, N. E. (2012). High-frequency neural activity and human cognition: Past, present, and possible future of intracranial EEG research. Progress in Neurobiology, 98, 279–301.

Lega, B., Jacobs, J., & Kahana, M. (2012). Human hippocampal theta oscillations and the formation of episodic memories. Hippocampus, 22(4), 748–761.

Long, N. M., Burke, J. F., & Kahana, M. J. (2014). Subsequent memory effect in intracranial and scalp EEG. NeuroImage, 84, 488–494. doi:10.1016/j.neuroimage.2013.08.052

Long, N. M., Sperling, M. R., Worrell, G. A., Davis, K. A., Gross, R. E., Lega, B. C., … Kahana, M. J. (2017). Contextually mediated spontaneous retrieval is specific to the hippocampus. Current Biology, 27, 1–6.

Manning, J. R., Jacobs, J., Fried, I., & Kahana, M. J. (2009). Broadband shifts in LFP power spectra are correlated with single-neuron spiking in humans. Journal of Neuroscience, 29(43), 13613–13620. doi:10.1523/JNEUROSCI.2041-09.2009

Merker, B. (2013). Cortical gamma oscillations: the functional key is activation, not cognition. Neuroscience & Biobehavioral Reviews, 37(3), 401–417.

Miller, K. J., Leuthardt, E. C., Schalk, G., Rao, R. P. N., Anderson, N. R., Moran, D. W., … Ojemann, J. G. (2007). Spectral changes in cortical surface potentials during motor movement. Journal of Neuroscience, 27, 2424–2432.

Miller, K. J., Weaver, K. E., & Ojemann, J. G. (2009). Direct electrophysiological measurement of human default network areas. Proceedings of the National Academy of Sciences, 106(29), 12174–12177.

Nunez, P. L., & Srinivasan, R. (2006). Electric fields of the brain. New York: Oxford University Press.

Osipova, D., Takashima, A., Oostenveld, R., Fernndez, G., Maris, E., & Jensen, O. (2006). Theta and gamma oscillations predict encoding and retrieval of declarative memory. J Neurosci, 26(28), 7523–7531. doi:10.1523/JNEUROSCI.1948-06.2006

Owen, A. M., Stern, C. E., Look, R. B., Tracey, I., Rosen, B. R., & Petrides, M. (1998). Functional organization of spatial and nonspatial working memory processing within the human lateral frontal cortex. Proceedings of the National Academy of Sciences, 95(13), 7721–7726.

Parsons, S., & Bynner, J. (2005). Does Numeracy Matter More? (Tech. Rep.). National Research and Development Centre for adult literacy and numeracy.

Sederberg, P. B., Schulze-Bonhage, A., Madsen, J. R., Bromfield, E. B., McCarthy, D. C., Brandt, A., … Kahana, M. J. (2007). Hippocampal and neocortical gamma oscillations predict memory formation in humans. Cerebral Cortex, 17(5), 1190–1196.

Solomon, E., Kragel, J., Sperling, M., Sharan, A., Worrell, G., Kucewicz, M., … Kahana, M. (2017). Widespread theta synchrony and high-frequency desynchronization underlies enhanced cognition. Nature Communications, 8(1), 1704. doi:10.1038/s41467-017-01763-2

Staudigl, T., & Hanslmayr, S. (2013). Theta oscillations at encoding mediate the context-dependent nature of human episodic memory. Current Biology, 23(12), 1101–1106.

Tulving, E. (1983). Elements of episodic memory. New York: Oxford.

Ueda, K., Brown, E. C., Kojima, K., Juhász, C., & Asano, E. (2015). Mapping mental calculation systems with electrocorticography. Clinical Neurophysiology, 126(1), 39–46. Retrieved from http://dx.doi.org/10.1016/j.clinph.2014.04.015 xdoi:10.1016/j.clinph.2014.04.015

Vansteensel, M. J., Bleichner, M. G., Freudenburg, Z. V., Hermes, D., Aarnoutse, E. J., Leijten, F. S. S., … Ramsey, N. F. (2014). Spatiotemporal characteristics of electrocortical brain activity during mental calculation. Human Brain Mapping, 35(12), 5903–5920. doi:10.1002/hbm.22593

Winawer, J., Kay, K. N., Foster, B. L., Rauschecker, A. M., Parvizi, J., & Wandell, B. A. (2013). Asynchronous broadband signals are the principal source of the bold response in human visual cortex. Current Biology, 23(13), 1145–1153.

Yushkevich, P. A., Pluta, J. B., Wang, H., Xie, L., Ding, S.-L., Gertje, E. C., … Wolk, D. A. (2015). Automated volumetry and regional thickness analysis of hippocampal subfields and medial temporal cortical structures in mild cognitive impairment. Human Brain Mapping, 36(1), 258–287.

